# The duration of nutrient limiting conditions can contribute to shaping subsequent diatom community composition; insights from laboratory experiments

**DOI:** 10.1101/2025.09.22.677915

**Authors:** Drajad S. Seto, Lee Karp-Boss, Mark L. Wells

**Affiliations:** Department of Oceanography, School of Marine Sciences, University of Maine, Orono, ME, USA; Department of Cell and Molecular Biology, College of the Environment and Life Sciences, University of Rhode Island, Kingston, RI, USA; Climate Change Institute, University of Maine, Orono, Mem USA

## Abstract

Climate-driven increases in global surface water temperatures are enhancing upper ocean stratification likely resulting in more prolonged periods of nutrient limitation. Although nutrient limitation in diatoms and their growth responses to increasing temperatures have been studied extensively, much less is known about their growth response to nutrient injection after prolonged durations of nutrient limitation. This study examines the growth response of three bloom-forming diatom species: *Pseudo-nitzschia pungens, P. australis*, and *Skeletonema costatum* after short-term (∼2 week) and prolonged (∼4 week) periods of nutrient limitation at five temperatures (9, 12, 15, 20, and 25°C). *Pseudo-nitzschia* species showed shorter lag times and higher growth rates than *S. costatum* after prolonged nutrient stress. These findings demonstrate that certain diatom species can exhibit faster growth recovery after prolonged nutrient limitation and in warmer conditions compared to others, providing new insights on drivers that shape phytoplankton communities.

## Introduction

Marine diatoms often dominate phytoplankton biomass in coastal and other high productivity regions of the oceans, particularly during spring blooms and upwelling events (Lassiter et al., 2006; Martin et al., 2011). This is a highly diverse group of species that exhibit a wide range adaptive mechanisms for survival under short-term nutrient stress, including changes in photosynthetic efficiency (Liefer et al., 2018), alteration in lipid composition (Machado et al., 2016), adjustment of sinking speeds (Du Clos et al., 2021), and altered metabolic pathways (Lampe et al., 2019). When nutrient stress progresses over longer time scales some species shift their metabolism to form resting spores as a strategy for survival (McQuoid & Hobson, 1996; Montresor et al., 2013; Von Dassow & Montresor, 2011). This shift involves both metabolic and morphological changes that include the development of a thick silica coatings (McQuoid & Hobson, 1996). Cells emerge from these dormant stages and recover vegetative growth once conditions improve. Although not widely studied, formation of resting stage has been only observed in centric diatom species so far (e.g., McQuoid & Hobson, 1996; Pelusi et al., 2019; Wang et al., 2024)

Not all diatoms form spores to survive nutrient stress. Some species, including certain pennate diatoms, appear to down-regulate or pause their growth stages in a type of “hibernation” that is not yet understood (Montresor et al., 2013), while others may rely on nutrient storage or lower minimum quotas (Tilman et al., 1982). These divergent strategies suggest that growth responses of diatom to an injection of nutrients after prolonged starvation may differ, whereby species not substantially reorganizing their metabolic or morphological structures potentially could resume vegetative growth faster once conditions improve. The subsequent staggered lag times for re-initiating exponential growth might give an advantage to some species over others mediating succession during bloom formation. Although these community changes may be modulated by grazing (Landry & Calbet, 2004), this influence is typically delayed in the case of rapidly forming diatom blooms.

With continued ocean warming and the increase in the frequency, duration, and intensity of heat waves over the past decade (Oliver et al., 2018), understanding the role of prolonged periods of seasonal stratification and nutrient limitation in shaping phytoplankton communities in costal ecosystems becomes even more relevant. While many studies have examined the effects of nutrient limitation and co-limitation on the physiology and ecology of phytoplankton, there is very little information about how the history of limitation (i.e., its duration) affects the recovery potential of a given species. Diatoms of the genus *Pseudo-nitzschia* in the coastal upwelling system of the California current have been often observed to dominate when upwelling is preceded by prolonged periods of abnormally warm conditions (Clark et al., 2019; Du et al., 2016; McCabe et al., 2016), but the nutritional history of the cells that generated these blooms is unknown and generally very difficult to assess in the field. Here, we conducted a laboratory study to examine how three coastal diatom species respond to macronutrient additions after Short-term (∼12 days) and Prolonged (∼27 days) periods of limitation; two species of *Pseudo-nitzschia*, a pennate genus that is not known to form resting spores, and one species of the genus *Skeletonema*, a centric genus that has the potential to produce resting spores in response to nutrient limitation (McQuoid, 2002; Montresor et al., 2013). We predicted that growth recovery from prolonged nutrient stress will differ between the two genera, with the two *Pseudo-nitzschia* species having a shorter lag time than *S. costatum*. We further examined if responses vary as a function of growth temperature to better understand the combined effects of warmer temperatures and nutrient limitation that cells may experience under prolonged periods of warming.

## Materials and Methods

### Culture conditions

We used non-axenic cultures of three diatom species: toxigenic pennate diatom *Pseudo-nitzschia pungens* from the Gulf of Maine (EBB1, Peter Countway, Bigelow Laboratory, ME) and *P. australis* from coastal water off Washington State (Bryan D. Bill and Vera Trainer, NOAA), and a non-toxic centric diatom *Skeletonema costatum* strain (CCMP778; isolated from the Caribbean Sea in the North Atlantic). Cultures were grown in sterile autoclaved media that was prepared with filtered seawater (0.7 μm GF/F, Whatman™, Pittsburg, PA, USA) collected from Frenchman Bay (ME, USA) or the University of Maine Darling Marine Center dock (Walpole, ME, USA). Filtered seawater was enriched with 16 µmol L^-1^ NO_3_ ^-^, 16 µmol L^-1^ (Si(OH)_4_, and 3 µmol PO_4_^3-^, conditions that reflect the deep water nutrient concentrations of the Gulf of Maine (Townsend et al., 2010), along with 25% of L1 trace metals and vitamins concentrations (Guillard & Hargraves, 1993). Hereafter, we refer to the medium as “Gulf of Maine (GoM) media”. Cultures were maintained under cool white fluorescent light (Philips TLD 36W/840, YZ36RL25) with a 14:10 h light:dark cycle.

### Experimental designs

Two independent experiments were conducted, with triplicate treatments. Experiment 1 aimed to determine the growth response of *P. pungens, P. australis* and *S. costatum* to short-term and prolonged nutrient limitation at a common sea surface temperature during spring/summer in the Gulf of Maine (16°C). Experiment 2 examined these limitation periods across five temperatures (9°C, 12°C, 15°C, 20°C, 25°C), reflecting Gulf of Maine seasonal norms and projected 2100 conditions (IPCC 2019).

### Experiment 1-Effect of nutrient limitation duration at 16°C

Cultures of *P. pungens, P. australis*, and *S. costatum* were grown in semi-continuous batch mode in 175 ml culture tissue flasks with GoM media at 16 ± 1°C (Fig. 1). Cells were acclimated for ≥ 10 generations before starting the experiment. Cells were considered ‘acclimated’ when growth rates varied less than 10% in consecutive butch cultures.

**Figure 1.**
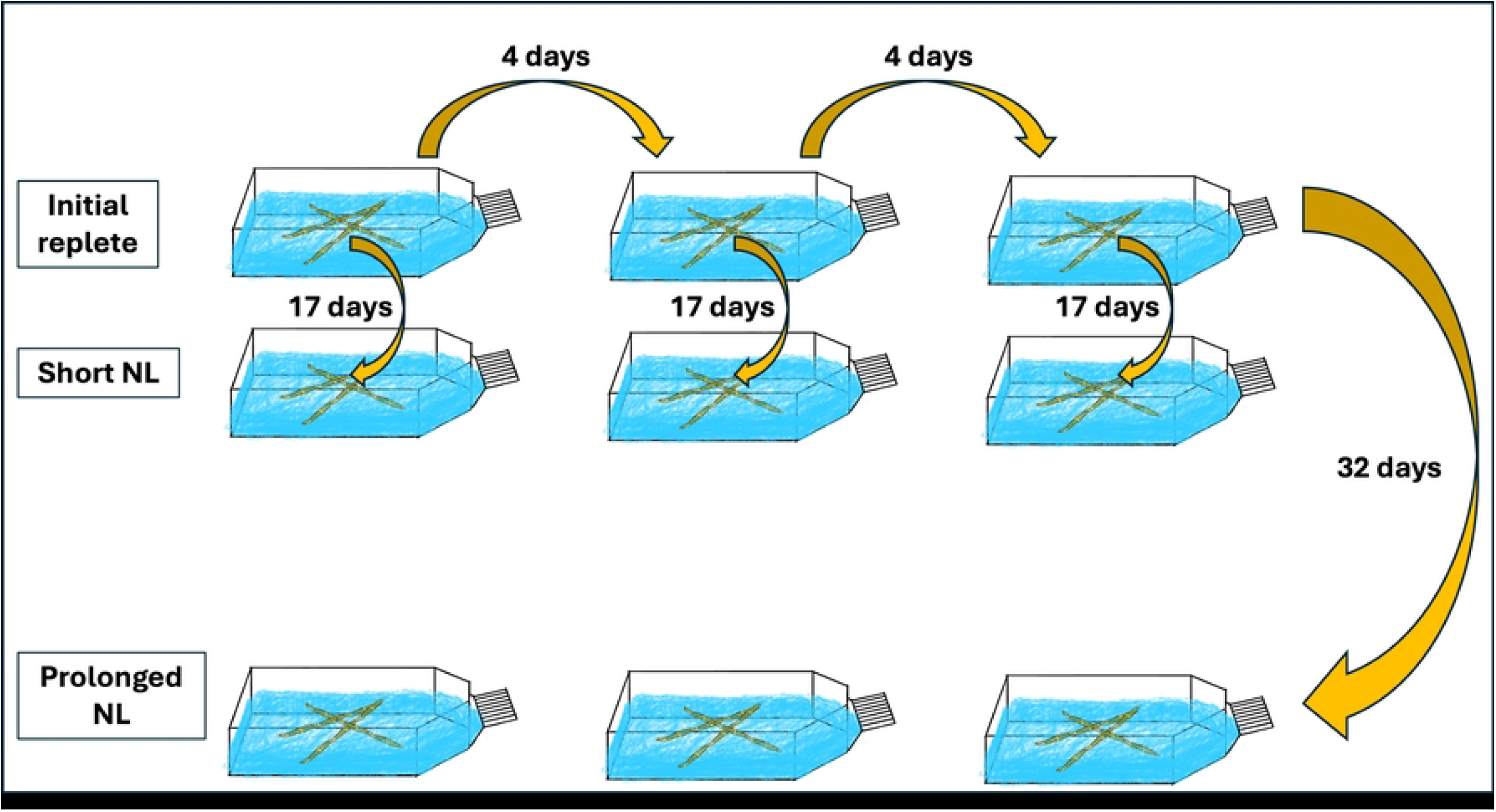
Experimental Design. Cultures were grown under three conditions: Initial replete, Short-term nutrient limitation (Short-term NL; 17 days total, with effective limitation for ∼12 days), and Prolonged nutrient limitation (Prolonged NL; 32 days total, with effective limitation for ∼27 days), all at ambient temperature. Arrows represent the time (in days) between transfers to fresh media

After acclimation, cells were transferred to fresh GoM media on Day 0 of the experiment, termed “Initial replete” serving as the growth reference for statistical comparisons with Short-term and Prolonged NL (Fig. 1). In pre-liminary experiments we determined that nutrient concentrations for *P. pungens* had decreased to 2.59 μM for nitrate and 1.54 μM for silicate by Day 4-5, while for *S. costatum*, the nutrient levels were 0 μM for nitrate and 0.92 μM for silicate (S4 Fig.). The growth of both species reached stationary phase on day ∼5. Cultures remained in stationary phase under the same light and temperature conditions for additional 12 days (i.e., a total of 17 days from the start of the experiment) to ensure nutrient depletion (hereafter, “Short-term nutrient limitation”) or 32 days (“Prolonged nutrient limitation”, ∼27 days post-depletion) (Fig. 1). A subset of these stationary cultures was transferred to fresh GOM media after these limitation periods and sub-samples were taken daily and preserved with 1% Lugol’s iodine for cell counts. Exponential growth rates were determined by cell counts using a Sedgwick-Rafter chamber for enumeration under inverted microscope (Nikon-TMS F). Growth experiments for *P. pungens* and *S. costatum* were conducted from July to September 2020; experiments with *P. australis* were conducted from May to July 2021; both experiments used the same basal seawater batch collected from the dock of the University of Maine Darling Marine Center.

### Experiment 2– Interactive effect of nutrient limitation duration and temperature

The same initial experimental plan was used for Experiment 2. Cultures were grown in triplicate 28 ml polycarbonate tubes (Nalgene™ Oak Ridge Centrifuge Tube) with 20 ml GoM media at 9°C, 12°C, 15°C, 20°C, and 25°C (Firstek, TG-5 model, Taiwan). After acclimation, cells were transferred at Day 0 and the experiment followed by the same protocol above. For Short-term nutrient limitation, 1 ml of each replicate was inoculated into fresh GoM media, and growth was tracked for ∼10 days. For Prolonged limitation, nutrients (16 µmol L^-1^ NO_3_^-^, 16 µmol L^-1^ (Si(OH)_4_, and 3 µmol PO_4_^3-^) and 25% of L1 trace metals and vitamins concentrations (Guillard & Hargraves, 1993) were added directly to initial cultures to avoid diluting cell abundances below detection. Because the multi-factorial nature of the experiment (with replications), manual cell counts were too time consuming and we used the faster chlorophyll fluorescence approach (Turner Design, USA, model 10-AU-005-CE) (Wood et al., 2005) to estimate growth rates, keeping in mind biases associated with this methods. We validated chlorophyll fluorescence measurements with cell counts for all species at 16°C across Initial replete, Short-term nutrient limitation (NL), and Prolonged NL conditions. Due to the exponential nature of cell growth and non-linear fluorescence responses, log 2-transformed cell counts were plotted against log2-transformed fluorescence units, yielding strong overall correlations (r^2^ = 0.97 for *P. pungens*, r^2^ = 0.88 for *S. costatum*, r^2^ = 0.87 for *P. australis*); condition-specific correlations are reported in S2 Table, with plots for the Initial replete, Short-term, and Prolonged NL in S2 Figure. Growth response experiments for *P. pungens* and *S. costatum* were conducted from December 2020 to February 2021 using Frenchman Bay filtered (GF/F) seawater as a basal media, while growth experiments with *P. australis* were conducted from May to July 2021 using filtered seawater from the dock of the University of Maine Darling Marine Center.

### Growth measurements

Specific growth rates (SGR) were calculated as:

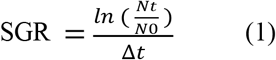

Where N_t_ and N_0_ are cell concentrations (Experiment 1) or fluorescence-derived biomass (Experiment 2) at time t and initial time, derived from exponential phase (Wood et al., 2005). For each replicate, N_0_ and N_t_ were measured individually. Lag times were defined as the number of days from nutrient resupply (Day 0) until the onset of the exponential growth phase, identified as the point where cell counts (Experiment 1) or in vivo fluorescence (Experiment 2) began a consistent logarithmic increase, determined by fitting an exponential growth model to daily measurements (Wood et al., 2005), if no exponential growth occurred within the experimental timeframe (e.g., ∼10 days), lag time was recorded as not applicable (NA).

### Statistical analysis

For Experiment 1, two-way ANOVA tested the effects of species (*P. pungens, P. australis, S. costatum*) and duration (Initial replete, Short-term, and Prolonged NL) on SGR and lag times, using raw triplicate data. Tukey’s HSD for post-hoc tests identified specific differences across all levels of duration: for SGR, species differences were examined within each duration (e.g., *P. pungens* vs. *S. costatum* in Initial replete, Short-term and Prolonged NLs) and duration differences within each species (e.g., Short-term NL vs. Initial replete for *P. pungens*); for lag times, duration differences were examined within each species (e.g., Short-term NL vs. Initial replete for *P. pungens*), and species differences were assessed within each species (e.g., *P. australis* vs. *P. pungens* in Prolonged).

For Experiment 2, two separate analyses were conducted using triplicate data: 1) temperature and 2) duration effects. For temperature effects, one-way ANOVA tested the effect of temperature (9°C, 12°C, 15°C, 20°C, 25°C) on SGR and lag times within each species and condition (Initial replete, Short-term and Prolonged NL). Tukey’s HSD post-hoc tests examined significant differences from 15°C (ambient/control) within each species and condition. For duration effects, two-way ANOVA tested the effects of species and duration within each temperature on SGR and lag times. Tukey’s HSD post-hoc testes examined significant differences from Initial replete conditions within each species and temperature. Analyses were performed in R version 2024.09.1+394.

## Results

### Experiment 1 - Effect of nutrient limitation duration at 16°C

Under Initial replete conditions, all species showed rapid growth with minimal lag times (Table 1, Figs. 2A&B). After exposure to Short-term NL, all species recovered upon nutrient resupply, with *P. pungens* maintaining a higher SGR than *S. costatum* (p < 0.01, Tukey’s HSD); *P. australis* showed intermediate SGR, significantly higher than *S. costatum* (p < 0.05). Lag times increased significantly for *P. pungens* and *S. costatum* (p < 0.05 vs. Initial replete) but remained unchanged for *P. australis* (p > 0.05). Following Prolonged NL, *P. pungens* and *P. australis* maintained a high SGRs compared to *S. costatum*, which showed no growth (p < 0.001) within five days period. Lag times further increased for *P. pungens* (p < 0.001 vs. Initial replete) and *P. australis* (not statistically tested vs. Initial replete due to insufficient replicates).

**Table 1.** Specific growth rates (SGR) and lag times for three diatom species under varying nutrient limitation durations at 16°C. NA indicates no growth observed. Lag time error is ± 1 day due to daily sampling. *P. australis* prolonged reflects n=1 and is excluded from species difference comparisons due to insufficient replicates. Means and standard deviation are calculated from triplicate data (except where noted). Asterisks (*) indicate significant changes in species differences (relative to Initial replete) across durations (p< 0.05, Tukey’s HSD). Dagger (†) indicate significant differences from Initial replete within each species (p< 0.05, Tukey’s HSD).

| Species | Condition | Temp. (°C) | SGR (Mean ± SD, d<sup>-1</sup>) | Lag Time (Mean ± SD, days) |
| --- | --- | --- | --- | --- |
| *P. pungens* | Initial replete | 16 | 1.58 ± 0.22 | 1 ± 0* |
| *P. pungens* | Short- NL | 16 | 1.24 ± 0.21*† | 2 ± 1† |
| *P. pungens* | Prolonged NL | 16 | 1.44 ± 0.1* | 3 ± 0† |
| *P. australis* | Initial replete | 16 | 1.38 ± 0.12 | 0 ± 1 |
| *P. australis* | Short- NL | 16 | $0.94 \pm 0.26^{\dagger}$ | $1 \pm 0$ |
| *P. australis* | Prolonged NL | 16 | 1.10 (n=1) | 7 (n=1) |
| *S. costatum* | Initial replete | 16 | $1.22 \pm 0.18$ | $1 \pm 0^*$ |
| *S. costatum* | Short- NL | 16 | $0.45 \pm 0.37^{\dagger}$ | $2 \pm 0^{\dagger}$ |
| *S. costatum* | Prolonged NL | 16 | $0^{\dagger}$ | NA |

**Figure 2.**
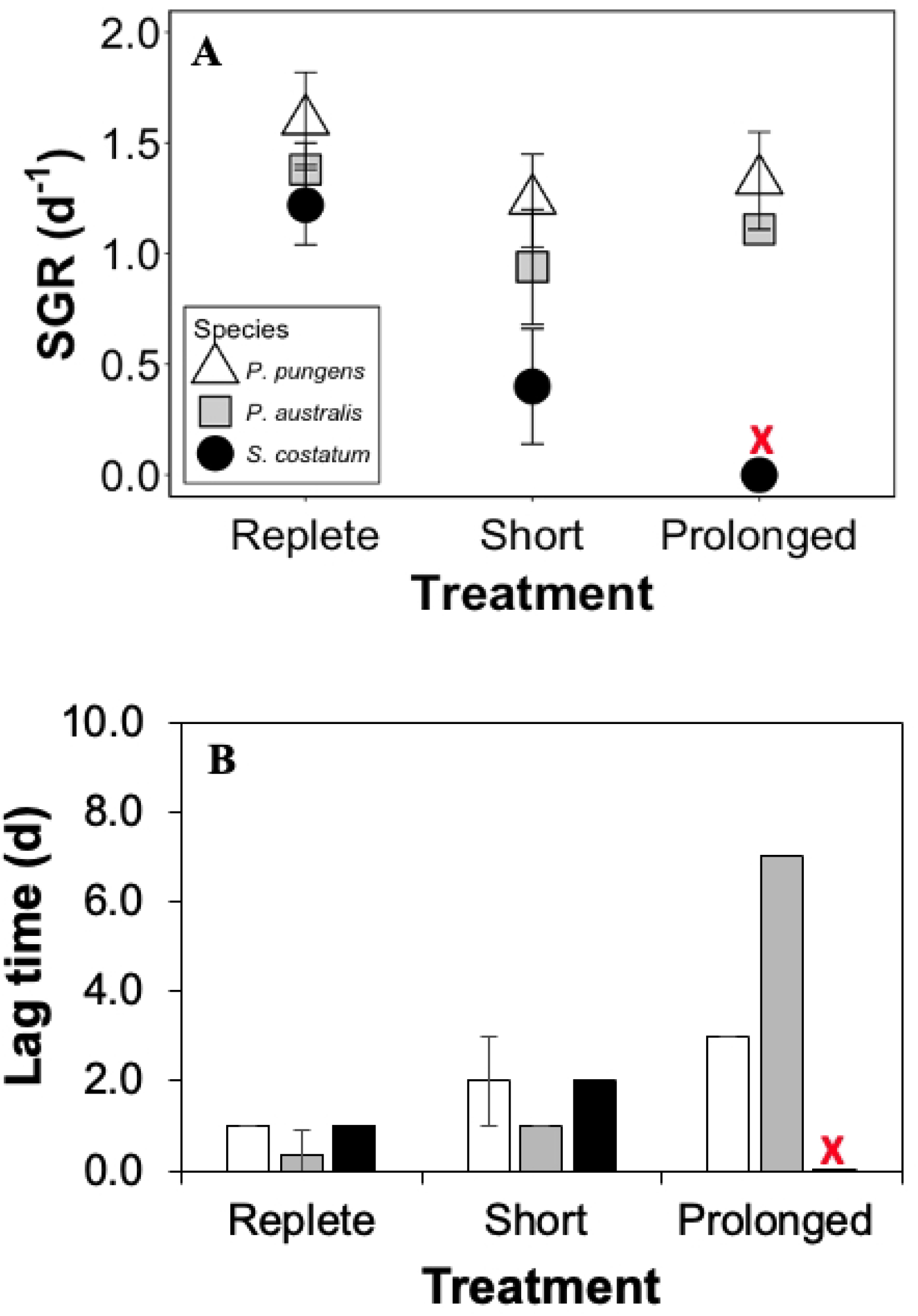
Experiment 1 results: Means and standard deviations of specific growth rates (SGR; panel A) and lag times (Panel B) of *P. pungens* (white triangle/bar), *P. australis* (grey square/bar), and *S. costatum* (black circle/bar) in the Initial replete nutrient treatment, and after Short-term and Prolonged nutrient limitation. Note that the *P. australis* treatment has one “replicate” (n=1) in the prolonged treatment.

Two-way ANOVA indicated significant effects of species (F(2,16) = 28.7, p < 0.001), limitation duration (F(2,16) = 34.2, p < 0.001), and their interaction (F(4,16)= 7.8, p < 0.01) on SGR (Fig. 2A). Tukey’s HSD showed that the specific growth rate of *P. pungens* was significantly higher than *S. costatum* in Short-term NL (p < 0.05) and Prolonged NL (p < 0.001) compared to Initial replete; *P. australis* in Prolonged NL (n=1) was not included in species difference comparisons due to insufficient replicates. Within species, Short-term NL SGR was significantly lower than Initial replete for *P. pungens* (p < 0.01), and *P. australis* (p < 0.05), and *S. costatum* (p < 0.01), and Prolonged NL SGR was lower than Initial replete for *S. costatum* (p < 0.001). For lag times (excluding *S. costatum* prolonged due to no growth, as lag time could not be measured), species (F(2,14) = 12.3, p < 0.001), duration (F(2,14) = 18.5, p < 0.001), and their interaction (F(3,14) = 4.2, p < 0.05) were significant (Fig. 2B). Tukey’s HSD showed that within *P. pungens* and *S. costatum*, Short-term NL (p < 0.05) and Prolonged NL (p < 0.001 for *P*.*pungens*) lag times were higher than Initial replete, while for *P. australis*, Short-term NL was not significantly higher (p > 0.05), and Prolonged (n=1) is reported descriptively; within Initial replete, lag time was lower than *P. pungens* and *S. costatum* (p < 0.05).

### Experiment 2 - Interactive effects of nutrient limitation duration and temperature

Under nutrient initial replete conditions, the SGR increased with temperature for all species, although the temperature optima varied (Fig. 3A, S1 Table). *Pseudo-nitzschia pungens* SGR increased from 0.36 ± 0.01 d^-1^ at 9°C to 1.40 ± 0.02 d^-1^ at 20°C (F(4,10) = 178.5, p < 0.001, one-way ANOVA), declining to 1.25 ± 0.10 d^-1^ at 25°C (p < 0.001 vs. 15°C), with no lag time across 9-25°C (Fig. 3B). *Pseudo-nitzschia australis* reached highest SGR at 15°C (1.60 ± 0.12 d^-1^, F(4,10) =115.6, p < 0.001), dropping to 0 d^-1^ at 25°C (p < 0.001 vs. 15°C), with a lag of 2 ± 1 days at 9°C (p < 0.001 vs 15°C). *Skeletonema costatum* SGR increased from 0.23 ± 0.03 d^-1^ at 9°C to 1.6 ± 0.12 d^-1^ at 25°C (F(4,10) = 144.8, p < 0.001), with a 2 ± 1 day lag at 9°C (p < 0.001 vs 15°C).

**Figure 3.**
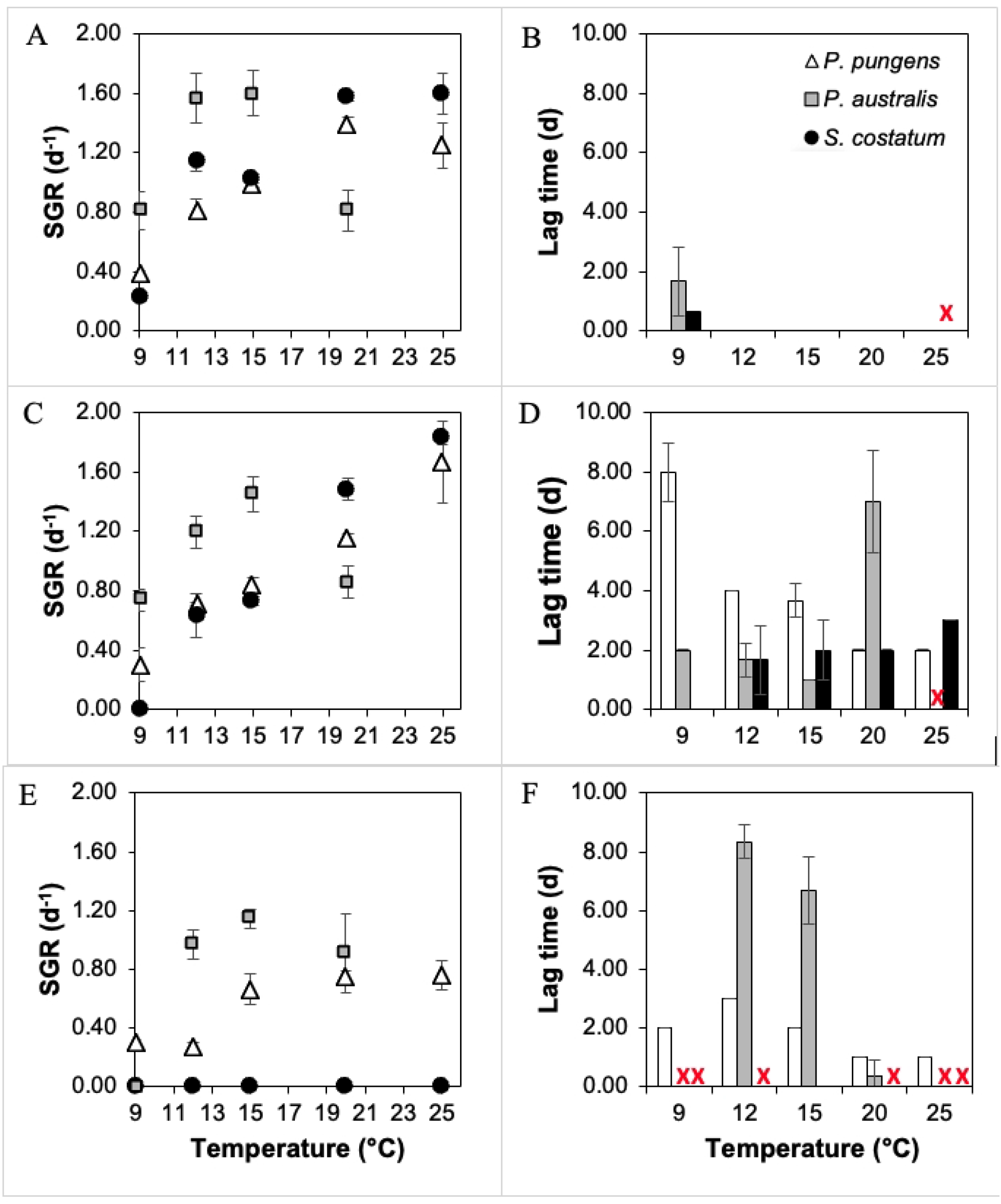
Experimental 2 results. Mean and standard deviations of specific growth rates (SGR) and lag times as a function of the temperatures of *P. pungens* (white triangle/bar), *P. australis* (grey square/bar), and *S. costatum* (black circle/bar) in Initial replete (A&B) (n=3), Short term NL (C&D) (n=3), and Prolonged NL (E&F) (n=3). Growth rates were determined from the regression slope on increasing in-vivo fluorescence during the exponential phase.

After Short-term NL, the SGR of *P. pungens* also increased from 0.71 ± 0.01 d^-1^ at 9°C to 1.67 ± 0.05 d^-1^ at 25°C (F(4.10) = 150.4, p < 0.001), with a longer lag at 9°C (8 ± 1 days, p < 0.001 vs 15°C, Fig. 3D). *Pseudo-nitzschia australis* peaked at 15°C (1.45 ± 0.11 d^-1^, F(4,10) = 149.2, p < 0.001), but showed no growth at 25°C (0.00 ± 0.00 d^-1^, p < 0.001 vs. 15°C) across all conditions, as cells failed to grow in the Initial replete conditions at this temperature (S1 Table); lag times increased to 7 ± 2 days at 20°C (p < 0.001 vs. 15°C). *Skeletonema costatum* did not grow at 9°C, peaking at 1.83 ± 0.03 d^-1^ at 25°C (F(4,10) = 187.6, p < 0.001), with lags of 2-3 days 9°C (Fig. 3C-D)

Growth responses of these species were markedly different after prolonged nutrient limitation. *Pseudo-nitzschia pungens* SGR increased from 0.29 ± 0.01 d^-1^ at 9°C to 0.76 ± 0.04 d^-1^ at 20-25°C (F(4,10) = 99.8, p < 0.001), with short lags of 1-3 days (Fig. 3E-F). *Pseudo-nitzschia australis* displayed no detectable growth at 9°C, peaking the SGR at 15°C (1.20 ± 0.12 d^-1^, F(4,10) = 87.4, p < 0.001), with longer lags of 8 ± 1 days at 12-15°C (p > 0.05, 12°C vs. 15°C), 1 ± 0 days at 20°C (p < 0.001 vs 15°C). As observed in Experiment 1, *Skeletonema costatum* did not respond well to prolonged nutrient limitation.

Within each temperature, treatment nutrient limitation duration significantly affected SGR and lag times (S1 Table). For example, at 15°C, SGR of *P. pungens* decreased in the Prolonged NL treatment (0.66 ± 0.10 d^-1^) compared to Initial replete (0.99 ± 0.02 d^-1^, p < 0.01) and increased *P. australis* lag time (8 ± 1 days) compared to Initial replete (0 ± 0 days, p < 0.001). At 20°C, Short-term NL increased *P. australis* lag time (7 ± 2 days) compared to Initial replete (0 ± 0 days, p < 0.001), while *S. costatum* SGR dropped to 0 d^-1^ in Prolonged NL compared to Initial replete (1.58 ± 0.07 d^-1^, p < 0.001). These patterns show the combined influence of nutrient limitation duration and temperature on diatom responses (S1 Table, Fig 3).

## Discussion

Increasing surface water temperatures have direct effects on cell metabolism and indirect effects from decreased nutrient availability as stratification intensifies (Behrenfeld et al., 2006; Li et al., 2020). While there is a rich literature on the direct effects of changing temperature and nutrient availability on diatoms (e.g., Fu et al., 2007; Padfield et al., 2016; Thomas et al., 2017), far less is known about how the duration of nutrient depletion associated with warming surface waters might influence diatom metabolism. We examine here whether the duration of nutrient stress may become a selective pressure influencing the composition of diatom communities, and how outcomes may vary under different growth temperatures.

The duration of nutrient limitation significantly impacted in the ability of these diatoms to recover after nutrient stress in our experiments. All three species recovered growth upon nutrient reinfusion after a short period of nutrient limitation (∼12 days) (Fig. 2A, 3C), though lag times varied with temperature (Fig. 2B, 3D). However, these responses diverged after prolonged (∼27 days) nutrient limitation. While both *Pseudo-nitzschia* species quickly responded to the nutrient additions, *S. costatum* was unable to recover its growth under these conditions over the duration of the experiment. Remarkably, lag times for *P. pungens* increased by only 1-3 days (relative to initial replete conditions) when nutrients were added after 27 days of depletion (Experiment 1 & 2; Figs. 2A & 3F). The resultant growth rates of both *Pseudo-nitzschia* species were somewhat lower relative to the initial replete conditions but still remained high (0.8 d^-1^ vs ∼1.5 d^-1^ for cells transferred to fresh media every 4 days; Table 1; Experiment 1, Fig. 2A). This rapid response was observed over the range of temperatures tested (Fig. 3E). The implication of these findings is that *S. costatum* would have a low probability of comprising a significant portion of the natural blooming phytoplankton assemblage after long periods of nutrient stress end through upwelling or enhanced mixing, whereas both *P. pungens* and *P. australis* likely could rapidly flourish under these conditions.

We cannot attribute a specific cause for these different growth responses in our experiments, as we could not visually confirm the presence of resting stages in the *S. costatum* within the experimental period. However, other internal cellular mechanisms also could have been at play. For example, the intracellular levels of sterol sulfates (Sts) associated with cell senescence increase as *Skeletonema* cells age, linked to an apoptosis-like death mechanism (Gallo et al., 2017). Alternatively, or perhaps in conjunction, nutrient stress of the genus *Skeletonema* has been shown to result in the over-production of reactive oxygen species (ROS), causing oxidative damage to cellular components (Wang et al., 2020). Regardless of the specific mechanism, the results demonstrate that the *S. costatum* strain tested here is poorly adapted for a rapid shift to the growth phase after prolonged nutrient stress when all other conditions are suitable for rapid growth.

The effect of temperature on this response also varied among the three diatom species but in a different way. Both *Pseudo-nitzschia pungens* and *S. costatum* increased their specific growth rates with increasing temperature, consistent with previous studies (e.g., Kim et al., 2015; Li et al., 2021). Short-term nutrient limitation did not alter this pattern (Figs. 3A&C). After prolonged nutrient stress *P. pungens* exhibited a rapid growth response but specific growth rates decreased by up to ∼50% at all temperatures relative to the initial replete conditions (Fig. 3E).

So, while the lag times were short, prolonged nutrient depletion still impeded the metabolic functioning of *P. pungens* to some degree.

In contrast to both *P. pungens* and *S. costatum*, specific growth rates of *P. australis* decreased at the highest temperature (25°C) (Fig. 3A,C,E), in agreement with previous findings (e.g., Clark et al., 2021; McCabe et al., 2016). Even so, its specific growth rate after prolonged nutrient limitation at 20°C was nearly identical to that under the initial replete conditions, and its lag time was even shorter (Figs. 3E&F) relative to short-term nutrient limitation (Figs. 3C&D). In other words, the findings indicate that this strain of *P. australis* can flourish after prolonged nutrient depletion at ≤20°C; i.e., ambient temperatures in most temperate coastal and offshore upwelling regimes.

While nutrient concentrations in cultures were not measure routinely in this study, our preliminary experiments confirm that nitrate was entirely depleted in *S. costatum* cultures by day 5. Similarly, nitrate declined in the *P. pungens* cultures from 19.11 µM at inoculation to 2.59 µM by day 5, while cell abundance increased from 333 to 18,083 cells mL^-1^. This corresponds to a drawdown of 16.52 µM nitrate, which, when divided by the increase in cell number (1.775×10^7^ cells L^-1^), implies ∼0.93 pmol N assimilated per cell. This estimate is comparable to reported quotas for smaller *Pseudo-nitzschia* species such as *P. subcurvata* (0.27-0.38 pmol N cell^-1^ (Zhu et al., 2017)) and since *P. pungens* typically has a larger cell size compared to *P. subcurvata*, its nitrogen quota are likely correspondingly higher. Given that only ∼2.6 µM nitrate remained by Day 5, nitrate limitation would have occurred shortly afterwards, as confirmed by the co-occurrence of senescence.

One could argue that factors other than nutrients, such as accumulation of toxic metabolic byproducts (e.g., oxylipins) or shift s in the associated microbial community, could have led to the senescence of cultures, as is often observed in F/2 or L1 media where growth reach senescence before nutrients are fully depleted. Because the initial nutrient concentration in the GoM media were significantly lower that F/2 or L1, and because cultures transferred to nutrient replete media on Day 17 (or 32) would also be diluted, such transfer potentially may have relieved stressors other than nutrient limitation. However, in the Experiment 2 under the Prolonged treatment, nutrients were directly injected to culture flasks (no transfer and dilution), so the observed responses represent the release from nutrient limitation and not the removal of secondary stressors. The consistent patterns observed from both Experiment 1 (at 16°C) and Experiment 2 (with the 15°C incubation), despite slight differences in methodology (transfer vs injection of nutrients) support that nutrient limitation in these cultures was essentially complete shortly after Day 5 of the experiments.

Further studies are needed to develop a mechanistic understanding of the underlying metabolic processes during short-term and prolonged periods of nutrient limitations across a broader range of taxa. Nevertheless, the findings from this study provide initial support for the idea that the duration of nutrient limitation can play a role in shaping phytoplankton communities after re-supply of nutrients, a factor that is not currently considered in models such as the phytoplankton Darwin model (e.g., Dutkiewicz et al., 2020; Fiksen et al., 2013). The implication is that the development and progression of bloom assemblages may be strongly influenced across a broader timeline of co-interacting bottom up, beginning far before bloom initiation.

In the absence of selective grazing pressures, taxa that are ready to resume growth immediately upon nutrient re-supply may be able to temporarily escape grazing control to initially dominate the community. Notably, field observations indicate that prolonged warming events are linked to the onset of *Pseudo-nitzschia* blooms along the West Coast of the United States. Perhaps the best example is the massive *Pseudo-nitzschia* bloom along much of the Western coast of N. America in 2015 (McCabe et al., 2016; Ryan et al., 2017). Anomalously warm (nutrient-depleted) waters were advected into the coastal region (the “warm blob”) in three winter months prior to the onset of seasonal upwelling. This upwelling created an intense, spatially continuous nearly monospecific *Pseudo-nitzscha* bloom for much of the western shore of N. America (McCabe et al., 2016; Ryan et al., 2017). Furthermore, on the opposite coast, anomalously warm and drought summer conditions (i.e., low nutrient influx from riverine flow or vertical mixing) in the Gulf of Maine region during 2016 preceded the first recorded, and spatially extensive toxic *Pseudo-nitzschia* bloom. Unlike the 2015 bloom off the west coast, this bloom happened during the fall turnover in 2016, replacing the more diverse species composition that normally is observed (Clark et al., 2019). These natural blooms are consistent with results from our laboratory experiments, showing that members of the genus *Pseudo-nitzschia* can quickly resume growth even after experiencing nutrient limitation for about one month, and suggesting why anomalous warming and prolonged nutrient limitation can potentially modulate the structure of diatom assemblages in coastal waters.

## Conclusion

The findings here show the extraordinary ability of two *Pseudo-nitzschia* spp. to quickly enter exponential growth after a prolonged nutrient limitation, in contrast to the *Skeletonema* spp. tested here. Although only three diatom species were tested here due to the intensive effort required for these long duration experiments, and the laboratory setting differs in many ways from ocean waters, the results demonstrate that the dynamics of nutrient stress is potentially a vital driver regulating natural diatom assemblages during at least the early stages of bloom development. The implication is that it is important to consider the nutritional history of species when evaluating their fitness to a changing environment. Future research should study a broader range of co-occurring species to better understand the prevalence of this response to macronutrient limitation but also consider the effects of prolonged micro-nutrient limitation on bloom composition. It is noteworthy that *Pseudo-nitzschia* species dominated the phytoplankton response in all mesoscale iron-enrichment experiments in High Nitrate Low Chlorophyll (HNLC) regions (Boyd et al., 2005). Better understanding of the different responses among diatoms after prolonged nutrient stress might come from transcriptomic experiments to elucidate the underlying cellular mechanisms. This work introduces a conceptual framework for how ocean warming may affect the timing and potential occurrence of blooms dominated by specific taxa in coastal and oceanic waters.

## Acknowledgment

We thank Peter Countway, Vera Trainer, and Brian D. Bill for providing *Pseudonitzschia* cultures, Sydney Greenlee for molecular identification of our diatom samples, David Townsend and Maura Thomas for macronutrient analysis, as well as Zexi Mao and David Carter for their assistance in measuring the fluorescence of the cultures.

## Supporting Information

S1 Fig. Experimental design for nutrient limitation and temperature effect in Experiment 2

S2 Fig. Validation of fluorescence proxy using log2 cell count vs log2 fluorescence unit for three diatom species

S3 Fig. Microscopic images *P. pungens* and *S. costatum* after Prolonged NL at 16°C in Experiment 1

S4 Fig. Macronutrients drawdown for *P. pungens* and *S. costatum*

S1 Table. Specific growth rate and lag time of three diatom species under varying nutrient limitation durations in Experiment 2

S2 Table. Condition-specific correlation between log2 cell counts and log 2 fluorescence unit for three diatom species in Experiment 1

## References

Behrenfeld, M. J., O’Malley, R. T., Siegel, D. A., McClain, C. R., Sarmiento, J. L., Feldman, G. C., Milligan, A. J., Falkowski, P. G., Letelier, R. M., & Boss, E. S. (2006). Climate-driven trends in contemporary ocean productivity. Nature, 444(7120), 752–755.

Boyd, P. W., Strzepek, R., Takeda, S., Jackson, G., Wong, C., McKay, R. M., Law, C., Kiyosawa, H., Saito, H., & Sherry, N. (2005). The evolution and termination of an iron-induced mesoscale bloom in the northeast subarctic Pacific. Limnology and Oceanography, 50(6), 1872–1886.

Clark, S., Hubbard, K. A., Anderson, D. M., McGillicuddy Jr, D. J., Ralston, D. K., & Townsend, D. W. (2019). Pseudo-nitzschia bloom dynamics in the Gulf of Maine: 2012– 2016. Harmful algae, 88, 101656.

Clark, S., Hubbard, K. A., McGillicuddy Jr, D. J., Ralston, D. K., & Shankar, S. (2021). Investigating Pseudo-nitzschia australis introduction to the Gulf of Maine with observations and models. Continental shelf research, 228, 104493.

Du Clos, K. T., Karp-Boss, L., & Gemmell, B. J. (2021). Diatoms rapidly alter sinking behavior in response to changing nutrient concentrations. Limnology and Oceanography, 66(3), 892–900.

Du, X., Peterson, W., Fisher, J., Hunter, M., & Peterson, J. (2016). Initiation and development of a toxic and persistent Pseudo-nitzschia bloom off the Oregon coast in spring/summer 2015. PloS one, 11(10), e0163977.

Dutkiewicz, S., Cermeno, P., Jahn, O., Follows, M. J., Hickman, A. E., Taniguchi, D. A., & Ward, B. A. (2020). Dimensions of marine phytoplankton diversity. Biogeosciences, 17(3), 609–634.

Fiksen, Ø., Follows, M. J., & Aksnes, D. L. (2013). Trait-based models of nutrient uptake in microbes extend the Michaelis-Menten framework. Limnology and Oceanography, 58(1), 193–202.

Fu, F. X., Warner, M. E., Zhang, Y., Feng, Y., & Hutchins, D. A. (2007). Effects of Increased temperature and CO2 on photosynthesis, growth, and elemental ratios in marine Synechococcus and Prochlorococcus (cyanobacteria) 1. Journal of Phycology, 43(3), 485–496.

Gallo, C., d’Ippolito, G., Nuzzo, G., Sardo, A., & Fontana, A. (2017). Autoinhibitory sterol sulfates mediate programmed cell death in a bloom-forming marine diatom. Nature communications, 8(1), 1292.

Guillard, R., & Hargraves, P. (1993). Stichochrysis immobilis is a diatom, not a chrysophyte. Phycologia, 32(3), 234–236.

Kim, J. H., Park, B. S., Kim, J. H., Wang, P., & Han, M. S. (2015). Intraspecific diversity and distribution of the cosmopolitan species Pseudo-nitzschia pungens (Bacillariophyceae): morphology, genetics, and ecophysiology of the three clades. Journal of Phycology, 51(1), 159–172.

Lampe, R. H., Wang, S., Cassar, N., & Marchetti, A. (2019). Strategies among phytoplankton in response to alleviation of nutrient stress in a subtropical gyre. The ISME Journal, 13(12), 2984–2997.

Landry, M. R., & Calbet, A. (2004). Microzooplankton production in the oceans. Ices Journal of Marine Science, 61(4), 501–507.

Lassiter, A. M., Wilkerson, F. P., Dugdale, R. C., & Hogue, V. E. (2006). Phytoplankton assemblages in the CoOP-WEST coastal upwelling area. Deep Sea Research Part II: Topical Studies in Oceanography, 53(25-26), 3063-3077.

Li, G., Cheng, L., Zhu, J., Trenberth, K. E., Mann, M. E., & Abraham, J. P. (2020). Increasing ocean stratification over the past half-century. Nature Climate Change, 10(12), 1116–1123.

Li, H., Xu, T., Ma, J., Li, F., & Xu, J. (2021). Physiological responses of Skeletonema costatum to the interactions of seawater acidification and the combination of photoperiod and temperature. Biogeosciences, 18(4), 1439–1449.

Liefer, J. D., Garg, A., Campbell, D. A., Irwin, A. J., & Finkel, Z. V. (2018). Nitrogen starvation induces distinct photosynthetic responses and recovery dynamics in diatoms and prasinophytes. PloS one, 13(4), e0195705.

Machado, M., Bromke, M., Júnior, A. P. D., Vaz, M. G. M. V., Rosa, R. M., Vinson, C. C., Sabir, J. S., Rocha, D. I., Martins, M. A., & Araújo, W. L. (2016). Comprehensive metabolic reprograming in freshwater Nitzschia palea strains undergoing nitrogen starvation is likely associated with its ecological origin. Algal Research, 18, 116–126.

Martin, P., Lampitt, R. S., Perry, M. J., Sanders, R., Lee, C., & D’Asaro, E. (2011). Export and mesopelagic particle flux during a North Atlantic spring diatom bloom. Deep Sea Research Part I: Oceanographic Research Papers, 58(4), 338–349.

McCabe, R. M., Hickey, B. M., Kudela, R. M., Lefebvre, K. A., Adams, N. G., Bill, B. D., Gulland, F. M., Thomson, R. E., Cochlan, W. P., & Trainer, V. L. (2016). An unprecedented coastwide toxic algal bloom linked to anomalous ocean conditions. Geophysical Research Letters, 43(19), 10,366-310,376.

McQuoid, M. R. (2002). Pelagic and benthic environmental controls on the spatial distribution of a viable diatom propagule bank on the swedish west coast 1. Journal of Phycology, 38(5), 881–893.

McQuoid, M. R., & Hobson, L. A. (1996). Diatom resting stages. Journal of Phycology, 32(6), 889–902.

Montresor, M., Di Prisco, C., Sarno, D., Margiotta, F., & Zingone, A. (2013). Diversity and germination patterns of diatom resting stages at a coastal Mediterranean site. Marine Ecology Progress Series, 484, 79–95.

Oliver, E. C., Donat, M. G., Burrows, M. T., Moore, P. J., Smale, D. A., Alexander, L. V., Benthuysen, J. A., Feng, M., Sen Gupta, A., & Hobday, A. J. (2018). Longer and more frequent marine heatwaves over the past century. Nature communications, 9(1), 1324.

Padfield, D., Yvon-Durocher, G., Buckling, A., Jennings, S., & Yvon-Durocher, G. (2016). Rapid evolution of metabolic traits explains thermal adaptation in phytoplankton. Ecology letters, 19(2), 133–142.

Pelusi, A., Santelia, M. E., Benevenuto, G., Godhe, A., & Montresor, M. (2019). The diatom Chaetoceros socialis: spore formation and preservation. European journal of phycology, 1–10.

Ryan, J., Kudela, R., Birch, J., Blum, M., Bowers, H., Chavez, F., Doucette, G., Hayashi, K., Marin III, R., & Mikulski, C. (2017). Causality of an extreme harmful algal bloom in Monterey Bay, California, during the 2014–2016 northeast Pacific warm anomaly. Geophysical Research Letters, 44(11), 5571–5579.

Thomas, M. K., Aranguren-Gassis, M., Kremer, C. T., Gould, M. R., Anderson, K., Klausmeier, C. A., & Litchman, E. (2017). Temperature–nutrient interactions exacerbate sensitivity to warming in phytoplankton. Global change biology, 23(8), 3269–3280.

Tilman, D., Kilham, S. S., & Kilham, P. (1982). Phytoplankton community ecology: the role of limiting nutrients. Annual review of Ecology and Systematics, 13(1), 349–372.

Townsend, D. W., Rebuck, N. D., Thomas, M. A., Karp-Boss, L., & Gettings, R. M. (2010). A changing nutrient regime in the Gulf of Maine. Continental shelf research, 30(7), 820–832.

Von Dassow, P., & Montresor, M. (2011). Unveiling the mysteries of phytoplankton life cycles: patterns and opportunities behind complexity. Journal of Plankton Research, 33(1), 3–12.

Wang, G., Huang, L., Zhuang, S., Han, F., Huang, Q., Hao, M., Lin, G., Chen, L., Shen, B., & Li, F. (2024). Resting cell formation in the marine diatom Thalassiosira pseudonana. New Phytologist.

Wang, H., Chen, F., Mi, T., Liu, Q., Yu, Z., & Zhen, Y. (2020). Responses of marine diatom Skeletonema marinoi to nutrient deficiency: programmed cell death. Applied and environmental microbiology, 86(3), e02460–02419.

Wood, A. M., Everroad, R., & Wingard, L. (2005). Measuring growth rates in microalgal cultures. Algal culturing techniques, 18, 269–288.

Zhu, Z., Qu, P., Gale, J., Fu, F., & Hutchins, D. A. (2017). Individual and interactive effects of warming and CO 2 on Pseudo-nitzschia subcurvata and Phaeocystis antarctica, two dominant phytoplankton from the Ross Sea, Antarctica. Biogeosciences, 14(23), 52815295.

